# Dual role of an essential HtrA2/Omi protease in the human malaria parasite: maintenance of mitochondrial homeostasis and induction of apoptosis like cell death under cellular stress

**DOI:** 10.1101/2022.03.16.484580

**Authors:** Shweta Singh, Gaurav Datta, Shaifali Jain, Vandana Thakur, Azhar Muneer, Mohd Asad, Shakir Ali, Asif Mohmmed

## Abstract

HtrA family of serine protease is known to play role in mitochondrial homeostasis as well as in the programmed cell death. Mitochondrial homeostasis and metabolism are crucial for survival and propagation of malaria parasite within the host. Here we have functionally characterized a *Plasmodium falciparum* HtrA2 (*Pf*HtrA2) protein, which harbours trypsin like protease activity that can be inhibited by its specific inhibitor, ucf-101. Transgenic parasite line was generated, using HA-*glm*S C-terminal tagging approach, for localization as well as for inducible knockdown of *Pf*HtrA2. The *Pf*HtrA2 was localized in the parasite mitochondrion during asexual life cycle. Genetic ablation of *Pf*HtrA2 caused significant parasite growth inhibition, decreased replication of mtDNA, increased mitochondrial ROS production, caused mitochondrial fission/fragmentation, and hindered parasite development. However, ucf-101 treatment had no effect on the parasite growth, suggesting non-protease/chaperone role of *Pf*HtrA2 in the parasite. Under cellular stress conditions, inhibition of *Pf*HtrA2 by ucf-101 reduced activation of the caspase like protease as well as parasite cell death, suggesting involvement of protease activity of *Pf*HtrA2 in apoptosis like cell death in the parasite. Under these cellular stress conditions, the *Pf*HtrA2 gets processed but remains localized in the mitochondrion, suggesting that it acts within the mitochondrion by cleaving intra-mitochondrial substrate(s). This was further supported by trans-expression of *Pf*HtrA2 protease domain in the parasite cytosol, which was unable to induce any cell death in the parasite. Overall, we show specific role of *Pf*HtrA2 in maintaining mitochondrial homeostasis as well as in regulating the stress induced cell death.

## Introduction

Malaria remains to be one of the major life-threatening diseases, especially affecting children in tropical and subtropical regions. In 2020, about 241 million cases have been reported causing 627,000 deaths worldwide [1] [2]. Artemisinin based combination therapies are mainly responsible for decline in cases in last few years; however, there is rapid emergence of drug resistant parasite strains against contemporary treatments including artemisinin combination therapies. Therefore, there is an urgent need to identify new drug-targets and develop new antimalarials [3, 4]. Identification of key metabolic pathways in the parasite is a prerequisite to identify unique and specific drug targets. Two parasite organelles of prokaryotic origin, the apicoplast and the mitochondrion, are essential for parasite growth and segregation. Metabolic pathways in these organelles are shown to be potential drug targets to obstruct the parasite development and segregation [5–9]. Therefore, understanding novel metabolic pathways in these organelles and understanding their functional role in the parasite, may help us to design novel anti-malarial strategies.

Mitochondria are known to play diverse role in various cellular activities including energy transduction and cell-death. Any disruptions in mitochondrial function and its dynamics could be lethal for the cell. In addition, mitochondria produce free oxygen radicals (reactive oxygen species, ROS) as a result of intrinsic physiological pathways, which could be also harmful for stability and functioning of mitochondrial proteins. Therefore, cells have developed molecular mechanisms to cope with diverse challenges imposed on mitochondrial integrity. In this direction, various heat shock proteins and chaperones ensure proper folding of new proteins and help in refolding of damaged proteins to maintain their optimal performance [10]. HtrA2 (high-temperature requirement A2)/Omi protein is among those heat shock proteins, which are present in human mitochondria, and suggested to be important modulators of molecular quality control. This protein combines the advantage of functioning both as chaperone and protease, based on requirement in the cell [10]. HtrA2/Omi belongs to a family of unique serine proteases, HtrA, which were first reported in *E. coli* [11]. HtrA proteins are shown to be vital for survival of bacteria only at higher temperature [12]. Therefore, it is suggested that HtrA proteins originated as a result of proteostatic stress in the cell, where they play role in protein quality control by recognizing misfolded protein or severely damaged proteins and their degradation if refolding is not possible [13]. In humans, there are four paralogs of HtrA proteases, HtrA 1, 2, 3 and 4. The most well studied among these is HtrA2, which has been also shown to play an important role in apoptosis. Human HtrA2/Omi harbours an N-terminal mitochondrial targeting sequence and gets localised to the mitochondria [14][15] The HtrA2/Omi is shown to play major role in protein quality control and maintaining the cellular homeostasis. Combination of catalytic domain and at least one C-terminal PDZ domain allows HtrA2/Omi to rapidly and reversibly switch their actions according to the needs of the cell, which makes HtrA2/Omi as a major quality control factor [10]. Protein structure analyses have shown that the HtrA2/Omi proteins forms pyramid-shaped homotrimer with the N-termini at the top and the PDZ domains at the bottom; the catalytic triad located in an interior of the trimer, which gets buried in the tight interface between PDZ and protease domains to form a catalytically inactive complex [16]. The chaperone activity of HtrA2 is carried out by recognising the hydrophobic features of unfolded polypeptides. The PDZ domains mediate initial substrate binding and subsequent translocation to the inner chamber. It has been also shown that HtrA2 activation as a protease requires binding of an activating peptide with the hydrophobic groove of PDZ. The PDZ domains of HtrA2 display a high affinity for hydrophobic peptides having the sequence YYF(V) at the C-terminus. Upon ligand binding, the PDZ domain triggers conformational reorganizations that result in the formation of a functional and accessible protease active site. This model is supported by the observation that the proteolytic activity of HtrA2/Omi is significantly augmented in absence of its PDZ domain and by peptide binding [17].

The HtrA2/Omi activity is shown to be essential for cell survival. Mutation or deletion of HtrA2/Omi induces ROS (Reactive Oxygen Species) generation, mitochondrial stress and triggers cell death pathway events [18]. In neural cells, the loss of HtrA2 has been shown to result in accumulation of unfolded proteins in mitochondria, decreased mitochondrial respiration, increased ROS production and ultimately neuronal cell death, in a manner similar to Parkinson’s disease [19, 20]. HtrA2/Omi is shown to protect mitochondria from cellular stresses, as in case of other homologous stress-adaptive proteins DegP and DegS in bacteria [21;13]. HtrA2/Omi is also shown to play role in apoptotic and autophagic signalling cell death pathways. Studies have shown that under stress conditions the activated HtrA2/Omi protease induces cell death in both a caspase-dependent as well as caspase-independent manner. HtrA2/Omi normally reside in mitochondrial intermembrane space, however, during stress conditions, like oxidative stress or heat stress, it breaches the inner membrane and migrates into the mitochondrial matrix [22–24]. In mitochondrial matrix, HtrA2/Omi converts into mature protease form and starts its functions by regulating cell death pathways [24]. Similarly in human neutrophils, the serine protease activity of HtrA2/Omi is required for caspase-independent, non-classical cell death pathway, where the protease targets substrate(s) within the mitochondria [15]. It was also shown that under apoptotic signals, HtrA2/Omi along with cytochrome c is relocated from the mitochondria to the cytoplasm [25]. The activated HtrA2/Omi protease translocated to the cytosol act as a proapoptotic protein, which inhibits the IAPs (Inhibitors of Apoptosis) and initiate activation of cascade of caspases, leading to cell death [14, 25, 26, 27]. In a brain ischemia rats model, the activated HtrA2/Omi gets translocated from mitochondria to the cytosol, which induces neuronal death [28], whereas its inhibition by specific inhibitor, ucf-101, attenuated ischemia-induced activation of caspase-8 and caspase-3 [29] In other process, HtrA2/Omi activates Beclin by acting on Hax-1 and disabling its activity resulting in release of Beclin-1, a homolog of autophagy 6, which promotes autophagy in the cell [24].

In the present study, we have functionally characterized the homologue of trypsin like serine protease HtrA2 in *P. falciparum* (PlasmoDB gene ID:PF3D7_0812200), *Pf*HtrA2, with a view to understand its role in mitochondrial homeostasis and parasite survival. We utilized HA-g*lm*S ribozyme system to study the localization and transient knockdown of *Pf*HtrA2 during asexual stages of the parasite. *Pf*HtrA2 was localized in the mitochondrion of the parasite; transient knockdown of *Pf*HtrA2 confirmed its essentiality for parasite survival and its role in development of functional mitochondria. We also show that PDZ domain plays role in regulating/blocking the protease activity of *Pf*HtrA2. Further, we show that the protease activity of *Pf*HtrA2 is required for apoptosis like cell death in the parasite induced by cellular stress conditions, where the protease act within the mitochondrion likely by cleaving an intramitochondrial substrate(s). Overall, these results suggest specific and essential role of *Pf*HtrA2 in maintaining mitochondrial homeostasis and parasite survival.

## Results

### Endogenous tagging of *PfHtrA2* with C-terminal HA-*glm*S ribozyme-tag and localization in transgenic parasites

To study functional essentiality of *Pf*HtrA2 (Fig. 1A), conditional knock-down using endogenous tagging strategy was utilized. We used single cross-over based homologous recombination method for C-terminal tagging of the native *pfhtrA2* gene with HA-*glm*S ribozyme system [30], so that the fusion protein gets expressed under the control of native promoter (Fig. 1B). Integration of HA-*glm*S at C-terminus of *pfhtrA2* locus was confirmed by PCR based integration analysis (Fig. 1C). A clonal population of parasite having HA-*glm*S tagged *pfhtrA2* locus was obtained using serial dilution and clonal selection method. This clonally selected homogenous parasite line was then used for all further experiments. The expression of *Pf*HtrA2-HA fusion protein (∼43kDa) was confirmed specifically in transgenic parasite (Fig. 1D) by western blot analysis using anti-HA antibodies; this band was not detected in wild-type 3D7 parasites (Fig. 1D).

**Figure 1:**
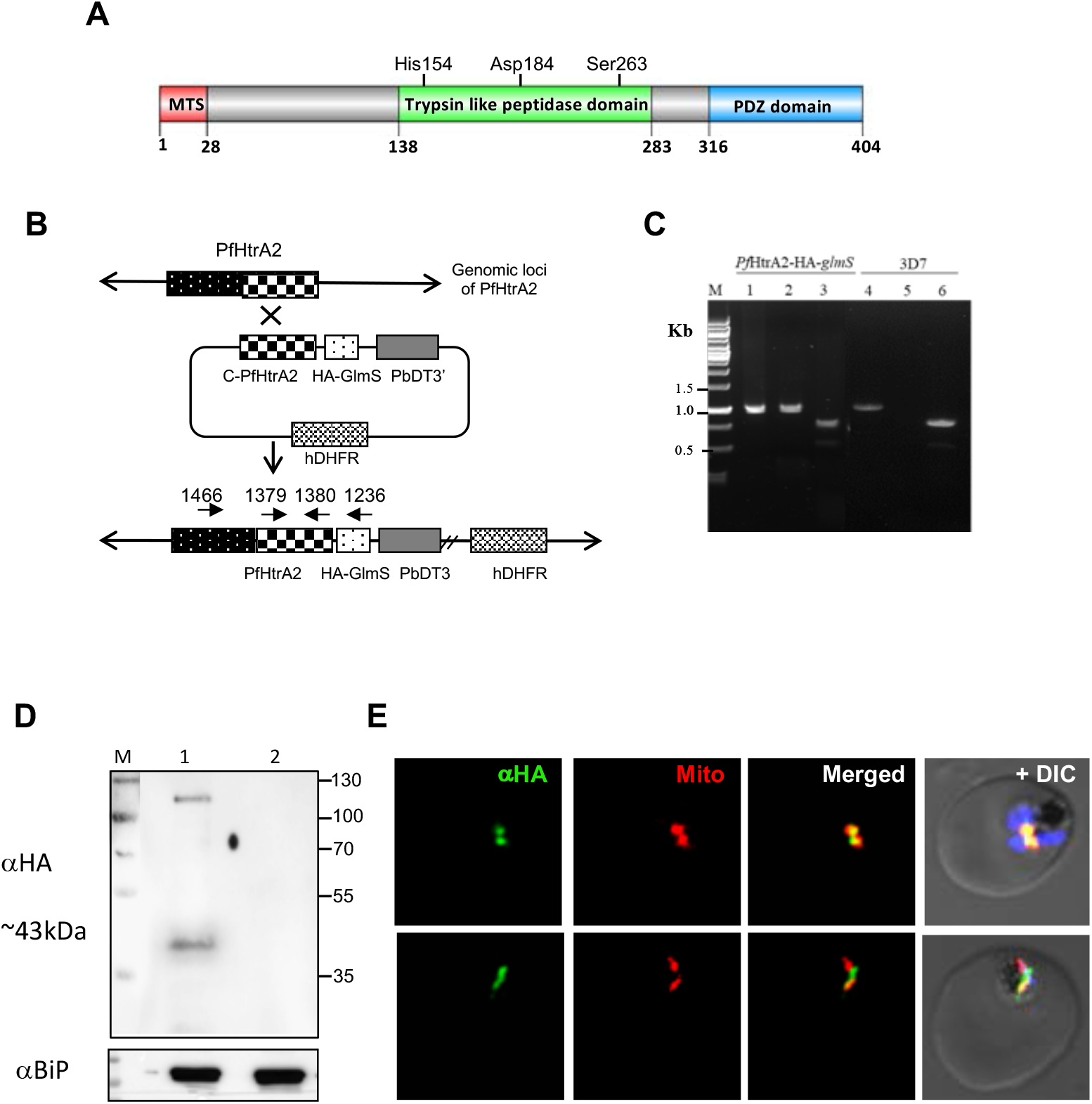
Generation of transgenic parasite expressing HA-*glm*S tag in the PfHtrA2 gene locus for localisation transient knock-down. (A) Schematic showing domain organisation of *Pf*HtrA2; the N-terminal mitochondrial targeting sequence, a trypsin-like protease domain, a C-terminal PDZ domain, and active sites residues of protease domain are marked. (B) Schematic diagram showing the strategy used to incorporate the HA-*glm*S at the 3’ end of endogenous locus of *pfHtrA2* through single cross over homologous recombination. (C) PCR -based analyses to confirm integration of plasmid in the target gene locus using total DNAs of the transgenic parasite culture (purified clonal parasite population) and wild type 3D7 parasite lines, locations of primers are marked in the schematic: lane 1 and 4 (primers1466A and 1380A) show amplification in both transgenic parasite with integrated plasmid and the wild-type line; lane 2 and 5 (primers1466A and 1236A) showing amplification only in the parasites with integration; lane 3 and 6 (primers 1379A and 1380A) show amplification in both integrant or wild type parasites. (D) Western blot analysis of lysate of transgenic and wild -type parasites using anti-HA antibody. The fusion protein band (∼43kDa) was detected in the transgenic parasites only (lane 1) and not in the wild-type parasites (lane 2). Blot ran in parallel with an equal amount of the same sample, probed with anti -BiP, was used as a loading control. (E) Fluorescent microscopic images of transgenic parasites immuno-stained with anti-HA antibody and labelled with MitoTracker. Fusion protein signal was observed in cellular organelle, mitochondrion, in transgenic parasites. The parasite nuclei were stained with DAPI and visualized by a confocal laser scanning microscope.

These transgenic parasites were studied for localization of the *Pf*HtrA2-HA fusion protein in asexual blood stages of the parasite by immuno-labelling and confocal microscopy. Anti-HA staining showed localization of fusion protein in a cellular organelle that showed characteristic shape, structure and division pattern of the parasite mitochondrion during the asexual blood-stage cycle. In young stages of the parasite, the mitochondrion is present as a small round-shaped structure close to the nucleus, in trophozoite-stage parasites the mitochondrion is present initially as elongated tubular structure and as branched structure in later stage (Fig. 1E). Indeed, staining by MitoTracker Red CMXRos showed overlap with the anti-HA labelling in these parasites (Fig. 1E). Further, co-localisation studies were also carried out using MitoTracker staining and immunolabelling with anti-*Pf*HtrA2 antibodies. The MitoTracker staining overlapped with anti-*Pf*HtrA2 labelling (Fig. S3). Thus, these results confirm that *Pf*HtrA2 is expressed in blood stage parasites and localises to the mitochondria in the parasites.

### Transient knock-down of *Pf*HtrA2 inhibits parasite growth, disrupts intra-erythrocytic parasite cycle and affects mitochondria development

To understand the functional significance of *Pf*HtrA2 in parasite survival and its possible role in mitochondrion development, we utilized the transgenic *Pf*HtrA2-*glm*S parasite line for inducible knock-down of *Pf*HtrA2 expression. The *glm*S-ribozyme gets activated in presence of glucosamine (GlcN) to cleave itself, which in turn leads to degradation of the associated *Pf*HtrA2-mRNA. Transgenic parasites treated with different GlcN concentrations (0mM, 2.5mM and 5mM) were analysed for selective knock-down of *Pf*HtrA2 protein. Treated parasites showed GlcN concentration dependent reduction in *Pf*HtrA2 levels (70-90%) (Fig. 2A). To avoid any potential GlcN-mediated toxicity observed at higher GlcN concentrations, all further experiments were carried out using 2.5mM of GlcN. To study the effect of the inducible knock-down of *Pf*HtrA2 (*Pf*HtrA2-iKD) on parasite growth, total parasitaemia was determined at different time points for three growth cycles (48h, 96h, 144h) after GlcN treatment. The *Pf*HtrA2-iKD set showed ∼80% inhibition of parasite growth compared to control set (Fig. 2B).

**Figure 2:**
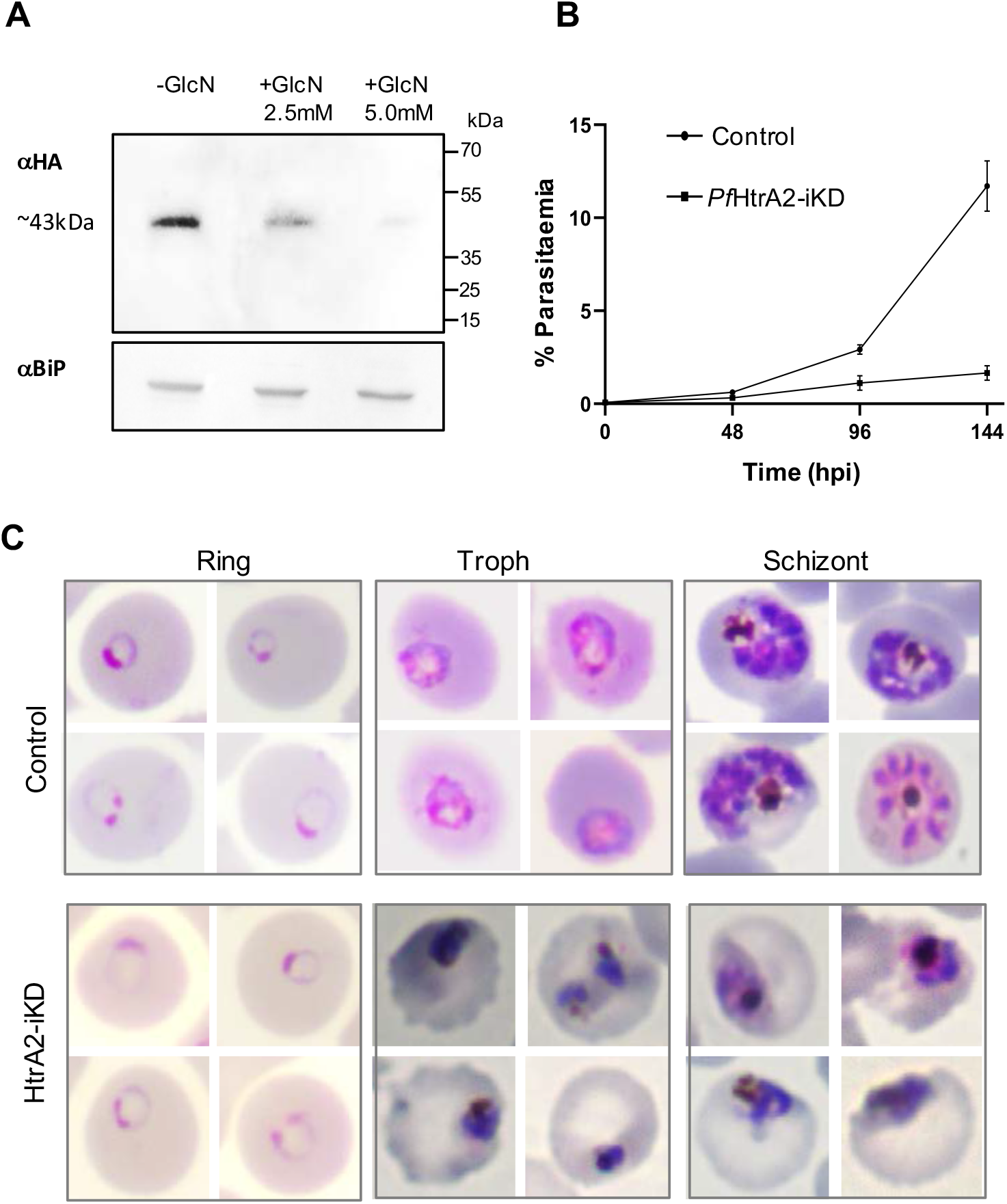
Inducible knock-down of PfHtrA2 protein in the transgenic parasites and its effect on growth and development asexual cycle of the parasites. (**A**) Western blot analysis using anti-HA antibody showing a reduction in the *Pf*HtrA2-HA fusion protein levels in transgenic parasites grown in presence of 2.5 mM glucosamine (+GlcN) as compared to control (-GlcN); a blot ran in parallel with an equal amount of the same sample was probed with anti-Bip antibodies as a loading control. (**B**) Graph showing *Pf*HtrA2-HA-*glm*S parasite growth in presence of 2.5 mM glucosamine (*Pf*HtrA2-iKD) as compared to control. Tightly synchronized ring-stage parasite culture of transgenic parasites grown with or without glucosamine (control and *Pf*HtrA2-iKD, respectively), and their growth was monitored as the formation of new rings determined at 48, 96 and 144hpi. All analyses were performed in triplicate or more (*n*=3); the error bars indicate standard deviations. (**C**) Giemsa-stained images of parasites showing effect on parasite morphology at different time points (0-120hpi) in parasite culture from control and *Pf*HtrA2-iKD sets.

The effect of selective knock-down of *Pf*HtrA2 on parasite development and morphology was assessed at different time points (Fig. 2C). As envisaged, in the control set, during each intra-erythrocytic cycle, parasites developed from ring to trophozoites to mature schizonts, and subsequently merozoites were released from these schizonts, which invaded new erythrocytes; the intra-erythrocytic cycle effectively increased the total parasitaemia for about 5-6-fold in every 48h cell-cycle under the culture conditions used in the study (Fig. 2B). In the *Pf*HtrA2-iKD set, parasite could only grow two-three folds in each cycle resulting in about 80% growth inhibition as compared to control set (Fig. 2B). Morphological observations of parasite intracellular development showed aberrant development of trophozoites and late-trophozoite stages (30-36hpi) in *Pf*HtrA2-iKD set as compared to control set (Fig. 2C). A large number of trophozoites were observed as darkly stained, pyknotic stressed parasites in the *Pf*HtrA2-iKD set; these parasites were not able to develop into schizonts at later time points. In the *Pf*HtrA2-iKD set, at 48 hpi, a number of stressed parasites were observed as compared to control set which had fully formed schizont stages (Fig. 2C).

### Downregulation of *Pf*HtrA2 levels hinders growth and segregation of the mitochondrion

Given that the *Pf*HtrA2 is a mitochondrial protease, we studied the effect of downregulation of *Pf*HtrA2 levels in the parasite on development of its mitochondrion. The growth and morphology of parasite mitochondria, in control as well as in the *Pf*HtrA2-iKD set, were observed at different time points during the erythrocytic cycle using MitoTracker stain. As expected, the mitochondria in control set showed normal growth and development: in majority of parasites at trophozoite stage (24-32hpi) the mitochondria appeared as elongated structures; in late-trophozoite/early-schizont stage (36-42hpi) majority of parasites showed much elongated and/or branched mitochondria (Fig. 3A, B, C). In *Pf*HtrA2-iKD set, only ∼20% of parasites mitochondria were able to develop into branched structures at 36hpi. Majority of parasites (>80%) in *Pf*HtrA2-iKD set showed fragmented structure or diffused punctate staining of MitoTracker dye with no clear organelle structure (Fig. 3A, B, C).

**Figure 3:**
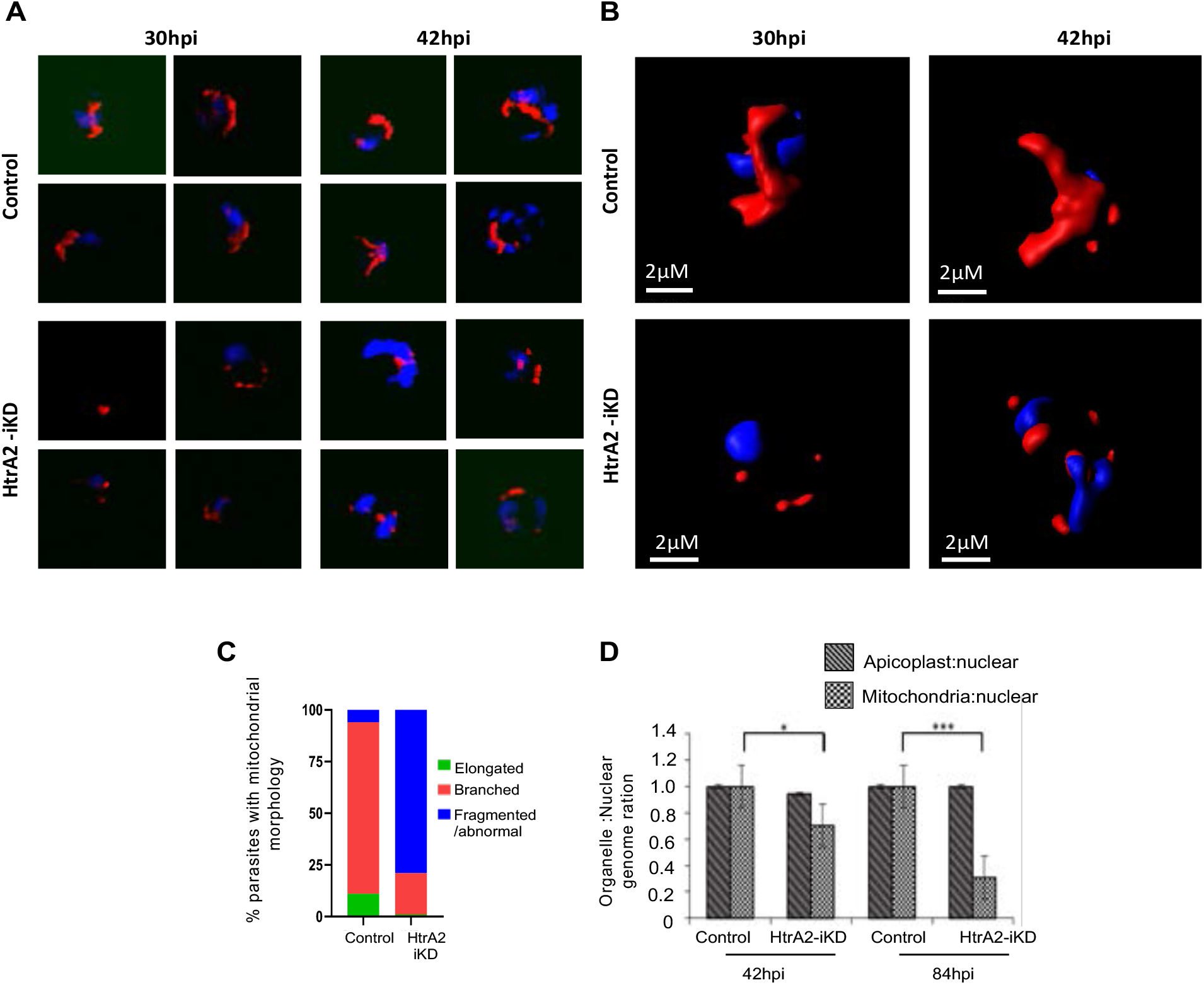
Effect of downregulation of *Pf*HtrA2 levels on growth and segregation of parasite mitochondrion. **(A)** Fluorescent microscopic images of MitoTracker stained *Pf*HtrA2-HA transgenic parasites at different time points (30-42 h) grown in presence (*Pf*HtrA2-iKD) or absence (control) of glucosamine. **(B)** Three-dimensional reconstruction using series of Z-stack images for selected parasite were made by IMARIS software showing intact mitochondria in the control parasites, whereas fragmented mitochondria in *Pf*HtrA2-iKD set. **(C)** Graph showing mitochondrial development profile in *Pf*HtrA2-iKD as compared to control set. Based upon mitochondrial morphology and staining pattern at 42hpi, three parasite groups were identified, and percentage parasites in each group are presented. **(D)** Quantitative PCR-based analyses using total DNAs isolated from *Pf*HtrA2-iKD parasites and control set during first and second cycle (36 hpi and 84 hpi); graph showing normalised genomic equivalents calculated for *tufA* present on apicoplast genome, and *cox-3* present on mitochondrial genome. Mitochondria to nuclear genome ratio was reduced in the *Pf*HtrA2-iKD set, while apicopolast to nuclear genome ratio remains unaffected.

To ascertain and quantitate the effect of iKD of *Pf*HtrA2 on mitochondria development and segregation in the parasite, we assessed replication of mitochondrial genome as compared to the replication of nuclear genome in the late trophozoite stages (42 hpi, 84 hpi) of the *Pf*HtrA2-iKD set. Quantitative PCR based analysis was carried out to estimate the genomic equivalents of the *cox-3* gene localised on the mitochondrial genome; in addition, genomic equivalence of *tufA* gene localised on the apicoplast genome was also determined. The mitochondrial genomic equivalence in the *Pf*HtrA2-iKD set showed two-fold reduction as compared to the control set within first growth cycle (Fig. 3D); these effects were pronounced in the second cycle. There was no significant effect on apicoplast genome as the genome equivalents for *tufA* were nearly similar in *Pf*HtrA2-iKD and control sets (Fig. 3D). These results clearly show that the down-regulation of *Pf*HtrA2 disrupts mitochondrial development and segregation.

### Downregulation of *Pf*HtrA2 disrupts mitochondrial function and induce oxidative stress

Given that knock-down of the HtrA2 in mammalian cells leads to loss of mitochondrial function and induce oxidative stress [18], we investigated effect of *Pf*HtrA2 knock-down on development of functional mitochondria and production of mitochondrial ROS in the parasite. The mitochondrial ROS production was assessed by using MitoSOX, which is an indicator for specific detection of mitochondrial superoxide in live cells. In control set, the trophozoite stage parasites showed faint mitochondrial staining using MitoSOX dye, whereas in the *Pf*HtrA2-iKD set the mitochondria showed significantly enhanced fluorescence suggesting higher levels of ROS (Fig. 4A). Quantitative levels of MitoSOX fluorescence in the mitochondrion for individual parasites were measured in the control and *Pf*HtrA2-iKD set, which showed significant increase of the mean normalised mean fluorescence intensity per parasite in the *Pf*HtrA2-iKD set (Fig. 4B). Further, the mitochondrial membrane potential was measured by staining with JC-1 dye. In the control set, the ratio of parasite population with mitochondrial red and cytosolic green staining was found to be ∼1.0 (Fig. 4C). However, in the *Pf*HtrA2-iKD set, there was a significant decline in the ratio of JC-1 red- and green-stained population to ∼0.5 (Fig. 4C), which clearly suggested loss of mitochondrial membrane potential.

**Figure 4:**
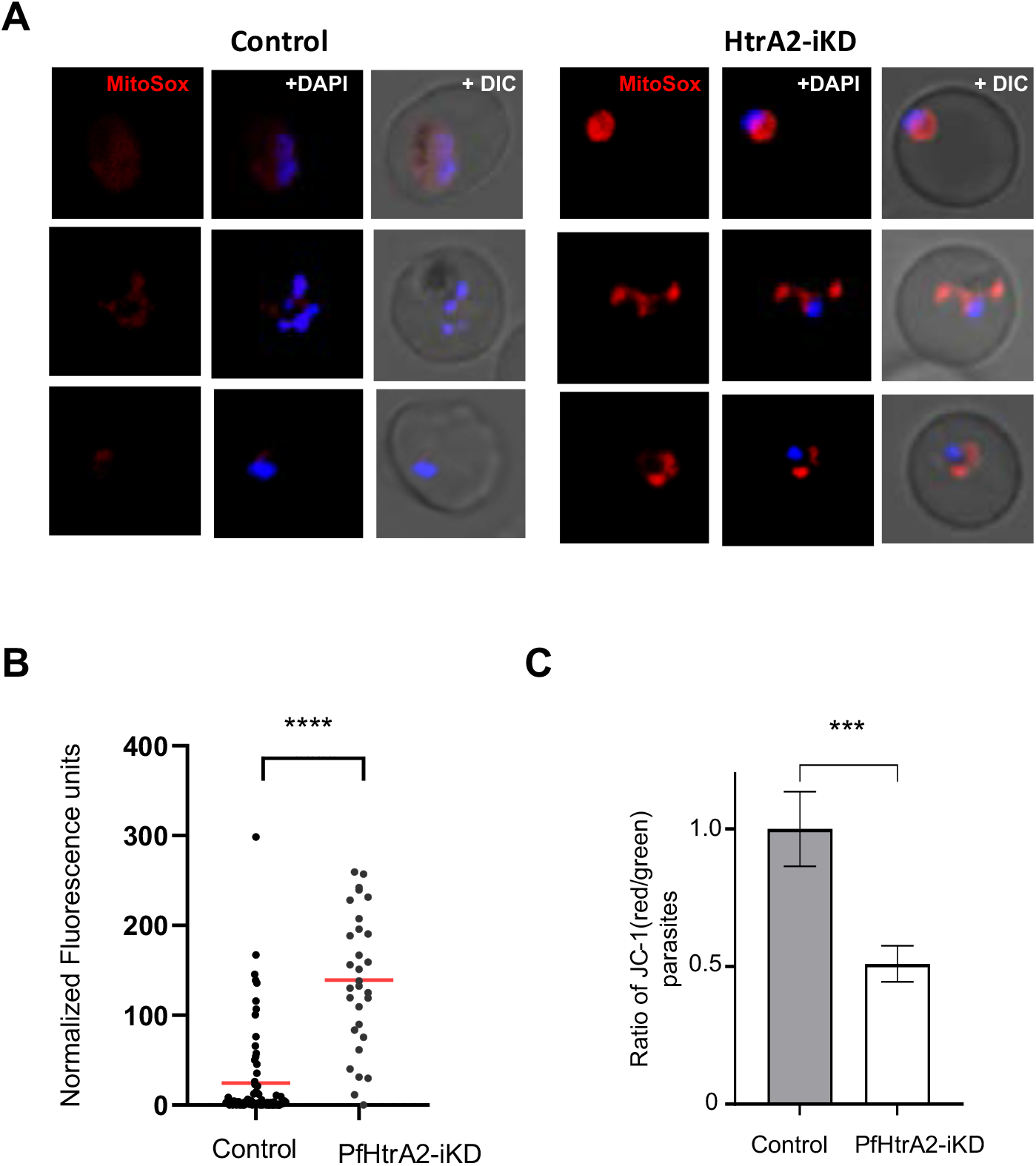
Effect of downregulation of *Pf*HtrA2 levels on mitochondrial oxidative stress and mitochondrial membrane potential. *Pf*HtrA2-HA tagged parasite were grown in the presence and in absence of glucosamine and stained with MitoSOX™ Red Mitochondrial Superoxide Indicator; enhanced red fluorescence of MitoSOX is detected in *Pf*HtrA2-iKD set suggesting higher levels of ROS. **(A)** Fluorescence microscopic images of parasites from *Pf*HtrA2-iKD set showing enhanced red fluorescence as compared to control. **(B)** Normalized fluorescence unit of in individual parasites in the *Pf*HtrA2-iKD set and control set (*n*=50), mean values are shown with horizontal bars with statistically significant differences (∗∗∗∗p < 0.001). **(C)** Graph showing reduction in ratio of JC-1 (red)/JC-1 (green) in parasite population after treatment with glucosamine (*Pf*HtrA2-iKD) as compared to control (*p < 0.05, **p < 0.01, ****p < 0.005 and ****p < 0.001).

### Characterization of *Pf*HtrA2 protease activity and identification of ucf-101 as its specific inhibitor

In order to characterise the protease activity of *Pf*HtrA2, the protease domain (135-290 aa; HtrA2-protease) was expressed as a histidine-tagged recombinant protein in *E. coli* and purified by affinity chromatography. The purified protein separated on SDS-PAGE (Fig. 5A; Fig. S4A-C) as a single band of ∼17 kDa, this protein was used to assess the protease activity using different small peptide substrates covalently linked to a fluorophore (AMC-amino-methyl-coumarin). These peptide substrates were specific for trypsin like protease activity (N-Succ-AFK-AMC), serine proteases (Z-LLVY-AMC, GGL-AMC), and cysteine proteases (ZFR-AMC, ZLR-AMC, and Z-LRGG-AMC). The *Pf*HtrA2-protease was able to significantly cleave the substrate specific for trypsin like protease (AFK-AMC) in a time dependent manner; however only weak activity was observed with any other substrate as compared to control wells (Fig. 4A, B). The kinetic analysis of recombinant *Pf*HtrA2-protease was determined by setting up reactions with varying concentrations of the substrate N-Succ-AFK-AMC (0-50 μM). The kinetic constant *K*_m_ was calculated to be 12.97 μM (Fig. S4D). The heterocyclic compound ucf-101 is a known inhibitor for the human HtrA2 protease activity (Cilenti et al. 2003). The inhibitory potential of ucf-101 was assessed in the standardized protease activity assay of *Pf*HtrA2. Ucf-101 was able to inhibit the protease activity of recombinant HtrA2-protease in a concentration dependent manner and the IC_50_ was calculated to be 8.1 μM (Fig. 5C).

**Figure 5:**
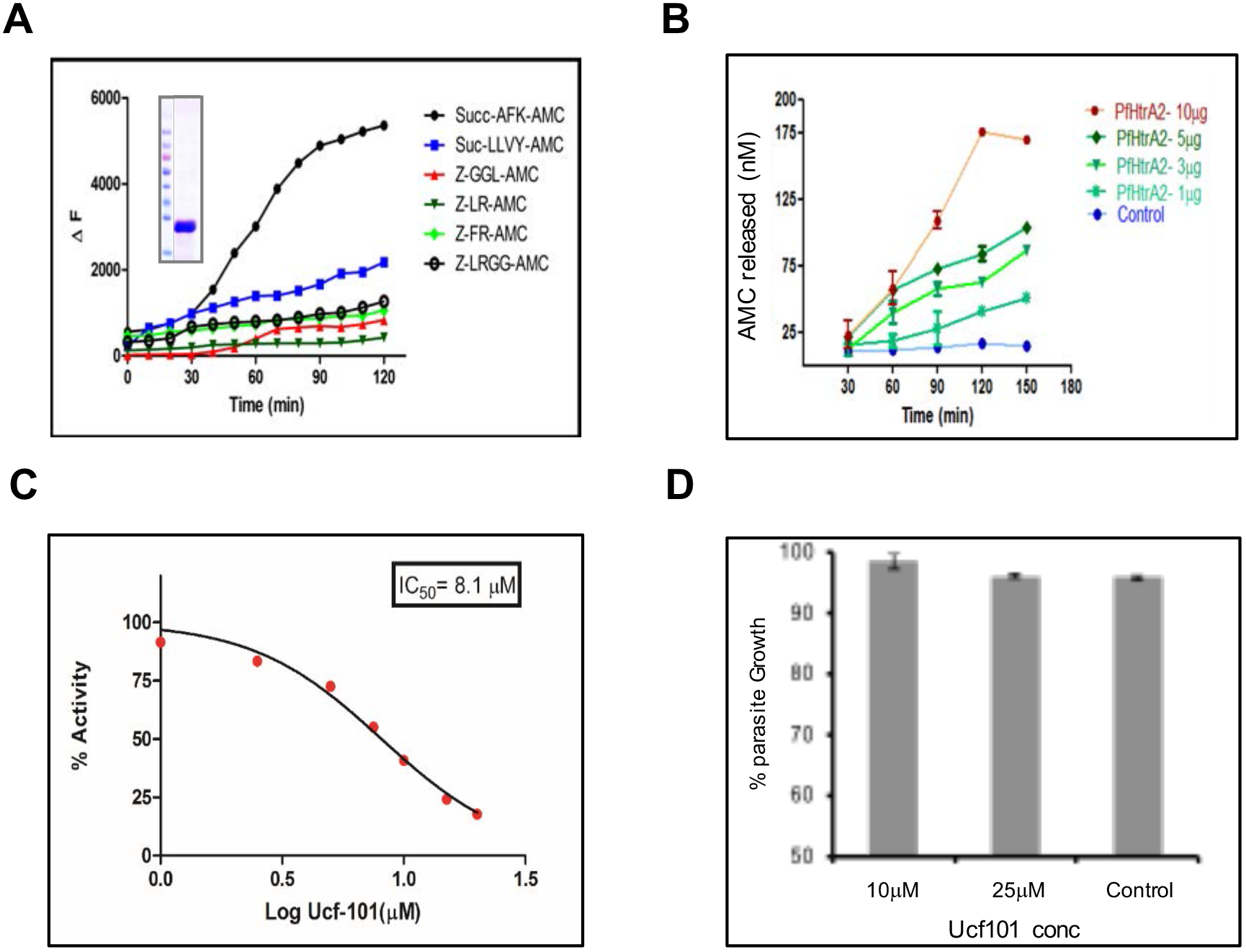
Protease activity assay for recombinant PfHtrA2. **(A)** Graphs showing activity of *Pf*HtrA2 with various fluorogenic peptide substrates. **(B)** Graphs showing activity of *Pf*HtrA2 using AFK-AMC substrate in a concentration dependent manner. **(C)** Graph showing concentration dependent inhibition of *Pf*HtrA2 activity by compound ucf101, the IC_50_ value was calculated to be 8.1μM. **(D)** Effect of various concentrations of ucf101 on parasite growth. The compound ucf101 showed no significant growth inhibition at different concentrations tested (> IC_50_ value on enzyme)

### *Pf*HtrA2 protease activity is not essential for parasite survival

The gene knock-down analysis showed that *Pf*HtrA2 protein is essential for parasite growth at the erythrocytic stages. To assess if the protease activity plays role in functional essentiality of the protein, we analysed parasiticidal efficacies of the *Pf*HtrA2 protease acitvity inhibitor ucf-101. The ucf-101 inhibitor is known to auto fluoresce at 543nm; treatment of asexual stage parasite cultures showed fluorescence in the parasite and specific accumulation in the parasite mitochondrion which overlapped with MitoTracker staining (Fig. S4E); which suggest that ucf-101 is membrane permeable and is able to reach at the site of action of *Pf*HtrA2 in the parasite. However, there was no effect on parasite growth for cultures treated with ucf-101 at concentration ∼IC_50_ and ∼IC_90_ for the HtrA2-protease activity (Fig. 5D). These results show that the protease activity of *Pf*HtrA2 is not essential for parasite growth and development.

### Interaction of PDZ-domain with protease-domain and inhibition of its activity

The PDZ domains are known to play role in protein-protein interaction and regulation of HtrA2 protease activity. To assess possible role of PDZ domain in maintaining the inactive HtrA2 protease under normal condition, we carried out protein-protein interaction analysis and suppression of protease activity by PDZ domain. The PDZ domain of *Pf*HtrA2 (316-404 aa) was expressed in with MBP-tag using *E. coli* expression system and purified with affinity chromatography as a ∼55kDa protein (Fig. 6A). Interaction of purified recombinant HtrA2-PDZ protein with recombinant HtrA2-protease (Fig. 6B), was confirmed by different strategies. In solution binding of both the recombinant proteins followed by immuno-pull down with antibodies against protease domain detected the HtrA2-PDZ in the eluates (Fig. 6C); in parallel assays, immuno-pulldown with antibodies against PDZ domain or with non-specific protein antibodies (anti-HDP) were used as positive and negative controls respectively. In another set of experiment, interaction of HtrA2-PDZ with HtrA2-protease was assessed on solid surface, followed by antibody-based detection; HtrA2-protease showed concentration dependent interaction with HtrA2-PDZ in this assay (Fig. 6D). A nonspecific recombinant *Pf*HDP protein was used as a negative control, which showed no significant interaction with HtrA2-PDZ.

**Figure 6:**
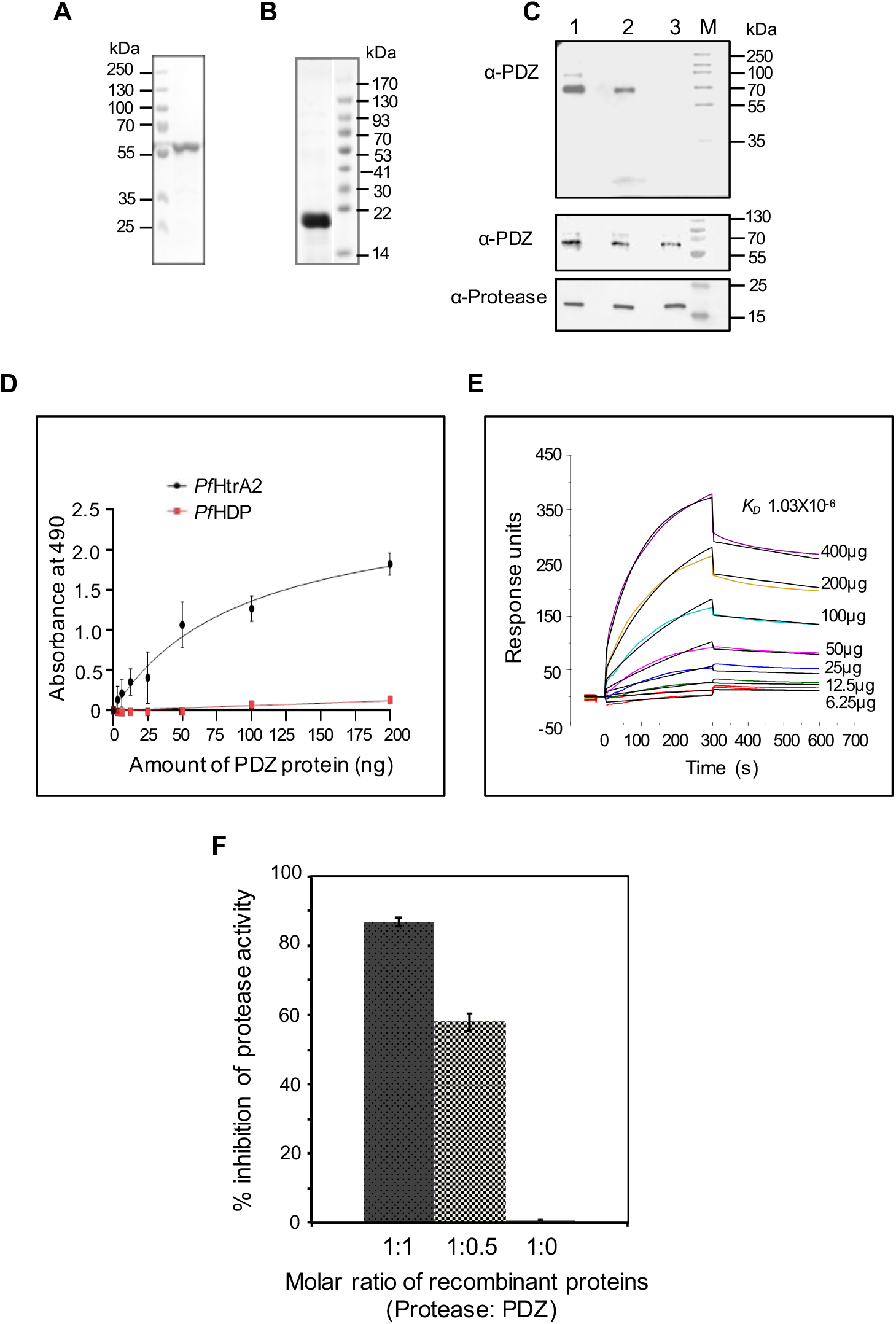
Interaction of protease and PDZ domains of *Pf*HtrA2, and its effect on its protease activity. **(A-B)** SDS-PAGE gels showing purified recombinant proteins *Pf*HtrA2-PDZ (A) and *Pf*HtrA2-protease (B). **(C)** Solution-binding assays of *Pf*HtrA2-PDZ and *Pf*HtrA2-protease recombinant proteins followed by *in vitro* pulldown of the protein complex using agarose beads labelled with anti-*Pf*HtrA2-PDZ (lane 1), anti-*Pf*HtrA2-protease (lane 2) or anti-*Pf*HDP (lane 3) antibody; presence of *Pf*HtrA2-PDZ protein in the pulled down protein complex was detected by western blot analysis using anti-PDZ antibody. Lower panel show western blot analysis to show presence of *Pf*HtrA2-PDZ as well as *Pf*HtrA2-protease recombinant proteins in the reaction mixture of each set. **(D)** *In vitro* solid-phase interaction of *Pf*HtrA2-protease recombinant proteins with increasing concentration of *Pf*HtrA2-PDZ; protein–protein interaction was determined by ELISA using anti-*Pf*HtrA2-PDZ antibody. **(E)** SPR based analysis of interaction of *Pf*HtrA2-protease proteins with *Pf*HtrA2-PDZ; global fitting of the sensograms obtained by flowing increasing concentrations (6.25-400 μg/ml) of *Pf*HtrA2-protease for 5 mins on *Pf*HtrA2-PDZ immobilized on the CM5 chip and dissociation for 5 mins under similar flow conditions. Buffer alone was used as control that showed no responses. **(G)** Interaction of *Pf*HtrA2-protease proteins with *Pf*HtrA2-PDZ suppresses its protease activity. Graph showing inhibition of *Pf*HtrA2-protease in presence of *Pf*HtrA2-PDZ; protease activity assay was carried out under different molar ratio of the two recombinant proteins, keeping the concentration of *Pf*HtrA2-protease constant.

Further, to ascertain the specificity of interaction and to quantitatively assess binding of HtrA2-PDZ with HtrA2-protease, a surface plasmon resonance-based biomolecular-interaction analysis was carried out. In this assay, the recombinant protein HtrA2-PDZ was immobilized permanently on sensor surface (CM5 chip) as ligand using EDC-NHS coupling, whereas the HtrA2-protease recombinant protein was injected in varying concentration and allowed to interact with immobilized HtrA2-PDZ. The assay channel showed typical association and dissociation curves as compared with the control cells, which showed little fluctuation due to solvent. The sensogram showed significant interaction of two recombinant proteins showing concentration dependent increase in binding of HtrA2-protease with immobilized HtrA2-PDZ (Fig. 6E). Kinetic analysis showed specific binding having strong equilibrium dissociation constant, *K_d_* value 1.03×10^-6,^ with the chi square value 32. However, no difference in response units was detected when HtrA2-protease was injected in either blank flow cells; recombinant *Pf*MSP1 was used as negative control in place of HtrA2-protease which showed no binding with immobilized HtrA2-PDZ protein. Overall, the data set obtained from these binding assays explicitly suggest the interaction of HtrA2-protease with HtrA2-PDZ. We further assessed any effect of this interaction of HtrA2-PDZ with HtrA2-protease on its activity; the HtrA2-protease activity assays were carried out in presence of recombinant HtrA2-PDZ. The HtrA2-PDZ was able to significantly inhibit the protease activity of HtrA2-protease (Fig. 6F); however, no inhibition was seen with a non-specific MBP fusion protein.

### Protein processing to release protease domain under cellular stress conditions

Homologues of HtrA2 are known to play a role in distinct cell death pathways, including caspase mediated cell death in eukaryotic cells. We have earlier shown the role of caspase like proteases under cellular stress conditions, including ER stress and mitochondrial stress, which induced apoptosis like cell-death in the parasites mediated by activation of caspase like cysteine proteases [7, 9]. Therefore, we assessed any role of *Pf*HtrA2 in ER-stress induced parasite cell death. We hypothesised that under ER stress, the full length *Pf*HtrA2 could undergo possible processing into a shorter fragment (43 and 25 kDa respectively, Fig 7A) and may translocate to the cytosol as shown in case of mammalian systems. Therefore, we first analysed possible processing of *Pf*HtrA2 (Fig. 7A) as well as any change in its localization in the parasites undergoing ER-stress induced cell-death. The synchronised parasite cultures under ER-stress at different time points were stained with MitoTracker, and immuno-stained with anti-*Pf*HtrA2 antibodies. The *Pf*HtrA2 was found to co-localise with MitoTracker staining after induction of the stress as in case of control set. Further, western blot analysis showed a band of full-length *Pf*HtrA2 protein (∼43 kDa) in control parasites, whereas for parasites under ER-stress, a processed form of *Pf*HtrA2 (∼25kDa) was detected (Fig. 7B, C).

**Figure 7:**
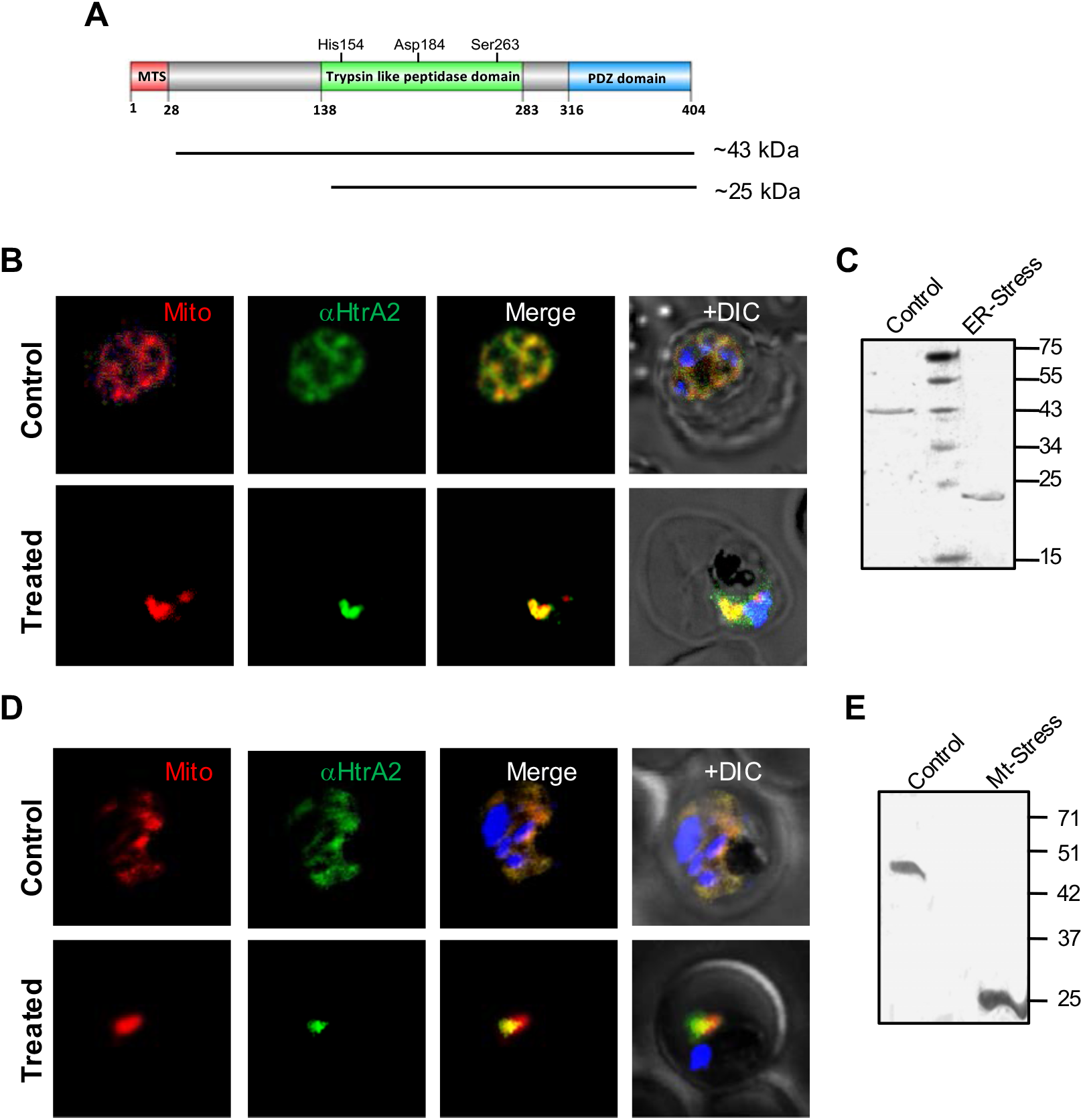
Localization and processing of *Pf*HtrA2 under cellular stress conditions. **(A)** Schematic diagram showing domain organization of *Pf*HtrA2 and indicating predicted sizes of full-length (∼43kDa) and processed form (∼25kDa). **(B)** Fluorescent microscopic images of parasites under ER-stress after immuno-labelling using anti-*Pf*HtrA2 antibody and staining with MitoTracker showing localization of *Pf*HtrA2 mainly in the mitochondria of the parasites as in the control set; parasite nuclei are stained with DAPI. **(C)** Western blot analysis of parasites under ER-stress showing presence of processed form *Pf*HtrA2 (∼25 kDa) as compared to full *Pf*HtrA2protease (∼43kDa) in the control set. **(D)** Fluorescent microscopic images of parasites under mitochondrial-stress after immuno-labelling using anti-*Pf*HtrA2 antibody and staining with MitoTracker showing localization of *Pf*HtrA2 mainly in the mitochondria of the parasites as in the control set; parasite nuclei are stained with DAPI. **(C)** Western blot analysis of parasites under mitochondrial-stress showing presence of processed form *Pf*HtrA2 (∼25 kDa) as compared to full *Pf*HtrA2protease (∼43kDa) in the control set.

To induce mitochondrial specific cellular stress, we utilized transgenic parasites expressing proteolytically inactive mutant ClpQ fused with FKBP degradation domain [*Pf*ClpQ (m)-DD], where regulated expression of mutant form of ClpQ protease induce dominant negative effect in the parasite leading to mitochondrial stress induced parasite cell death [8]. Localization studies in these transgenic parasites also showed *Pf*HtrA2 majorly remains to be localised in the mitochondria after induction of mitochondrial stress (Fig. 7D); further, western blot analysis of these parasites detected a ∼25 kDa band as compared to ∼43 kDa band of full length *Pf*HtrA2 protein, as in case of ER-stressed conditions (Fig. 7E).

### *Pf*HtrA2 protease activity plays role in cellular-stress induced apoptosis like cell-death

In mammalian cells, the HtrA2 follows different pathways to regulate activation of caspases, directly and indirectly [15, 31]. Since cellular stresses are shown to induce activation of caspase like cysteine proteases leading to apoptosis like cell-death in the parasite [7, 9], we assessed any possible role of *Pf*HtrA2 in this pathway. After induction of ER-stress a large percentage of parasites (∼50%) were found to be CaspACE positive, showing activation of caspase like cysteine proteases (Fig. 8A, S5A, C). The parasite cultures treated with ucf-101 (at IC_50_ concentration) under ER-stress condition, showed significant reduction in CaspACE positive parasites; in this set the CaspACE stained population was only 30%, suggesting ∼40% reduction in caspase activation as compared to the ER-stressed set without ucf-101. Similarly, when parasites were treated with caspase inhibitor z-VAD-FMK under ER-stress condition, there was significant reduction in CaspACE stained population (Fig. 8A). To further ascertain the role of *Pf*HtrA2 in caspase-like proteases mediated cell death in ER-stressed parasites, we assessed the effect of ucf-101 and z-VAD-FMK on cell death among these parasites by TUNEL assay, using the same experimental setup as described above. Inhibition of *Pf*HtrA2 by ucf-101 and inhibition of caspase-like proteases by z-VAD-FMK showed similar levels of effect on parasite cell-death in ER stressed parasites. Both the compounds showed ∼40% reduction in cell death under the ER-stress condition (Fig. 8B).

**Figure 8:**
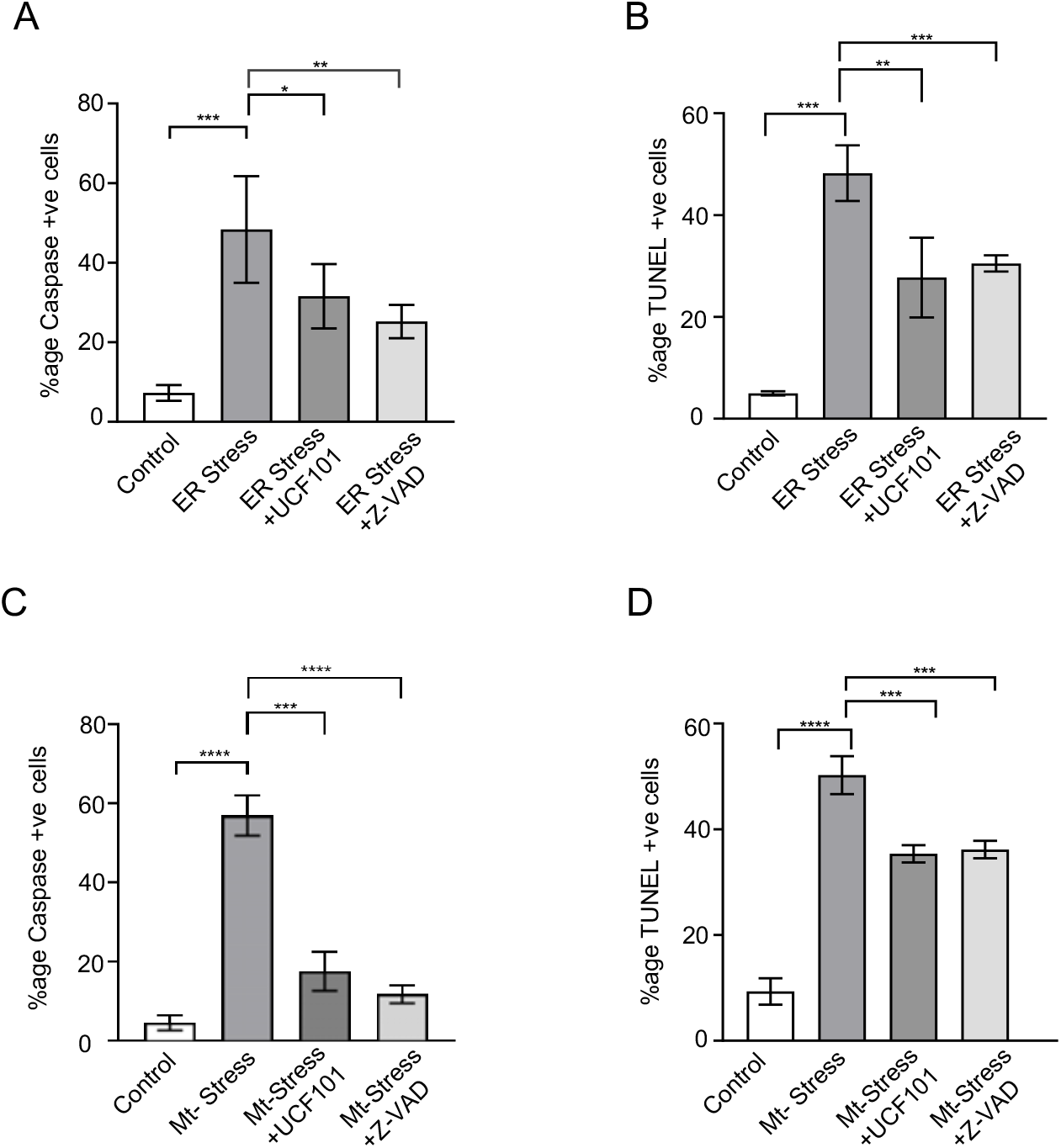
Inhibition of *Pf*HtrA2 protease activity reverses cellular stress induced activation of z-VAD-FMK binding caspase like proteases and parasite cell death. (**A**) Graph showing percentage of CaspACE-labelled parasites in the cultures after induction of ER-stress in presence or absence of inhibitor of *Pf*HtrA2 protease activity, ucf101, and in presence or absence of inhibitor of caspase like protease, z-VAD-FMK. (**B**) Graph showing percentage of TUNEL-labelled parasites in parallel sets for ER-stress (**C**) Graph showing percentage of CaspACE-labelled parasites in the cultures after induction of mitochondrial-stress in presence or absence of inhibitor of *Pf*HtrA2 protease activity, ucf101, and in presence or absence of inhibitor of caspase like protease, z-VAD-FMK. (**D**) Graph showing percentage of TUNEL-labelled parasites in parallel sets for mitochondrial stress. (*p < 0.05, **p < 0.01, ****p < 0.005 and ****p < 0.001).

We further analysed if *Pf*HtrA2 mediates activation of caspase like proteases in the mitochondrial stressed parasites. Induction of mitochondrial stress led to activation of caspase like cysteine proteases in a large percentage of parasites (∼60%) as shown by CaspACE staining (Fig. 8C, S5B, D). However, when parasites under mitochondrial stress were treated with ucf-101, there was a significant reduction in CaspACE stained population (∼60% reduction) as compared to mitochondrial stress alone (Fig. 8C). Treatment with z-VAD-FMK also resulted in ∼80% reduction in CaspACE staining (Fig. 8C). As in case of ER-stress, inhibition of *Pf*HtrA2 by ucf-101 and caspase-like proteases by z-VAD-FMK showed similar effect on mitochondrial stress induced cell-death. In both sets, there was 30-40% decrease in parasite cell-death under mitochondrial stress (Fig. 8D).

### Regulated trans-expression of *Pf*HtrA2 protease-domain and its effect on cell-death

We further ascertained that the processed and activated *Pf*HtrA2-protease plays role in initiating cell death cascade from within the mitochondria and not by getting released in the cytosol. For this, the protease domain of *Pf*HtrA2 was overexpressed as a transgene in the parasites. Since PDZ domain was found to inhibitory to protease activity (Fig. 6F), the PDZ domain was not included in the transgene. To avoid any deleterious effect of constitutive overexpression of the protease on the parasite, the protein levels in the parasite were regulated by tagging it with *E. coli* DHFR degradation domain (DDD) at the C-terminus along with the HA-tag. In addition, transgenic parasite line overexpressing active site mutant of *Pf*HtrA2-protease were also generated with same strategy to use as control. The parasite lines were labelled as *Pf*HtrA2-P-DDD and *Pf*HtrA2-P(mut)-DDD respectively (Fig. 9A). In these transgenic parasites, the DDD system enables the folate analogue trimethoprim (TMP) to stabilise the fusion protein, whereas removal of TMP results in degradation of the fusion protein by proteasome machinery. These transgenic parasites showed expression of *Pf*HtrA2-P-DDD and *Pf*HtrA2-P(mut)-DDD fusion protein of expected size (∼ 35 kDa) (Fig. 9B) in respective +TMP sets, which was not detected in absence of TMP. Further, the immunofluorescence assay using anti-HA antibodies also detected the fusion proteins in the parasite cytosol for both the transgenic parasite lines after treatment with TMP (Fig. 9C).

**Figure 9:**
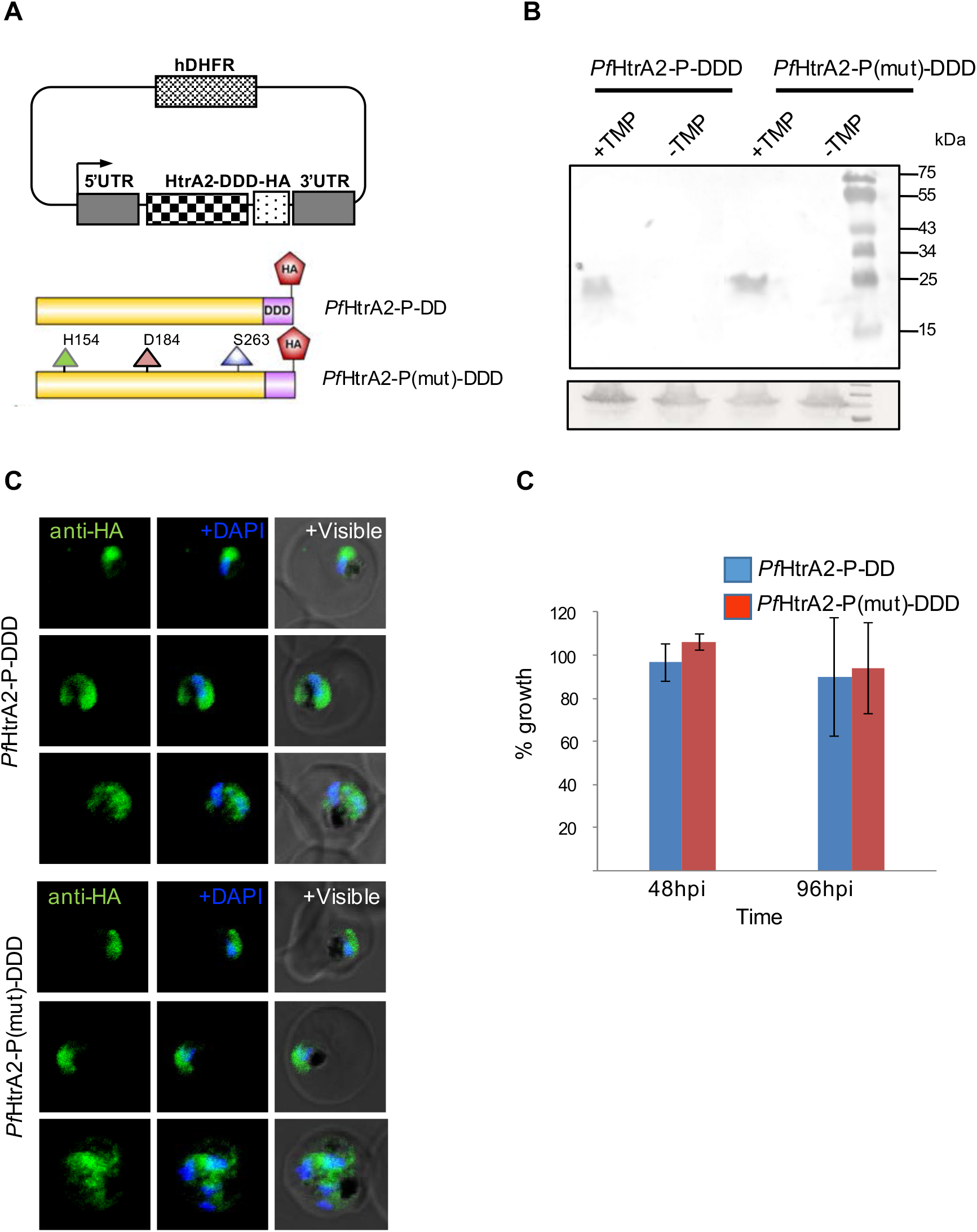
Trans-expression of *Pf*HtrA2-protease in parasite cytosol is not detrimental for the parasite. **(A)** Schematics of plasmid constructs and transgene for trans-expression of *Pf*HtrA2-protease and its mutant version in the parasite. The transgene corresponding to *Pf*HtrA2-protease and its triple mutant version (*Pf*HtrA2-P-DD and *Pf*HtrA2-P(mut)-DD) were expressed under the hsp86 promoter, whereas levels of these proteins are regulated by *E. coli* DHFR-degradation domain (DDD) tag. Expression of fusion proteins was induced in presence of trimethoprim (TMP). **(B)** Western blot analysis of lysate of transgenic parasites grown in presence (+TMP) or absence (-TMP) of trimethoprim, using anti-HA antibody. Expression of fusion protein ∼25 kDa, was not detected in +TMP set for both the parasite lines; a parallel blot probed with anti-BiP antibodies as loading control. **(C)** Fluorescent microscopic images of transgenic parasites grown in presence of trimethoprim, expressing *Pf*HtrA2-P-DD and *Pf*HtrA2-P(mut)-DD, and immune-stained with anti-HA antibody. The fusion proteins were present in the parasite cytosol in both cases. **(D)** Graph showing percentage growth of transgenic parasite cultures in presence of trimethoprim as compared to control set grown in absence of trimethoprim.

We next investigated effect of expression of *Pf*HtrA2-protease in the parasite cytosol on its growth. However, no effect on parasite growth was found in *Pf*HtrA2-P-DDD as compared to control set, similarly there was no significant difference in the growth of *Pf*HtrA2-P-DDD and *Pf*HtrA2-P(mut)-DDD parasites (Fig. 9D). Further, we assessed any induction of cell death by measuring CaspACE and TUNEL staining after induction of expression of transgene. However, no clear staining was observed in any of the two transgenic parasite lines; only background staining was detected as in case of control sets.

## Discussion

A number of metabolic pathways in the mitochondrion and the apicoplast, two parasite organelles of prokaryotic origin, are considered as potential drug targets in the parasite. HtrA2 is a serine protease which localizes in the mitochondrial membrane space, and is known to play role in mitochondrial protein quality control and homeostasis, as well as in regulation of autophagy and cell-death [10, 14, 32–36]. Here, we have identified a homologue of HtrA2 in human malaria parasite *P. falciparum* (*Pf*HtrA2) and carried out detailed localization, gene knock-down and cellular functional analyses to understand its role in mitochondrial homeostasis. The *Pf*HtrA2 protein comprise of an N-terminal mitochondrial targeting sequence (MTS), a trypsin-like protease domain, and a C-terminal PDZ domain, with conserved active site residues. Biochemical characterisation of the recombinant *Pf*HtrA2 protease domain confirmed that it is an active trypsin-like protease. Further, we confirmed localization of *Pf*HtrA2 in the parasite mitochondria, as predicted by the presence of mitochondrial targeting sequence.

To understand its functional role, we utilized transient gene-knock down strategy. A number of genetic tools have been developed for transient down regulation of target proteins in *Plasmodium* by tagging native gene [37]. Here we have tagged the native gene at the C-terminus with a ribozyme system [30] for its regulatable knock-down. Transient down regulation of *Pf*HtrA2 caused severe parasite growth inhibition and disruption in the development of trophozoites into schizonts stages; further, this also caused disruption of mitochondrial development in the parasite, combined with loss of mitochondrial membrane potential and induction of mitochondrial oxidative stress as evident by increase in mitochondrial ROS levels. Loss of HtrA2 in neuronal cells is shown to cause increased accumulation of unfolded components of the respiratory machinery in mitochondria, defects in the electron transport chain and enhanced production of ROS [20]; similarly, HtrA2 knockdown in embryonic fibroblast and human kidney cells is shown to disrupt the mitochondrial homeostasis leading to loss of mitochondrial membrane potential and increase in ROS production [38]. Our results for the first time suggest a key role of *Pf*HtrA2 in regulating mitochondrial homeostasis and development of functional mitochondrion in the parasite *P. falciparum*. HtrA2 has been shown to perform a dual function in the eukaryotic cell, in addition to acting as a mitochondrial chaperone, as described above, it can also act as a protease to regulate cellular proteins. In neural cells, loss of proteolytic function of HtrA2 results in accumulation of its direct substrate, p66Shc, which cause ROS generation and induction of apoptosis [39–41]. Although, direct substrate of *Pf*HtrA2 are not known in *Plasmodium*, but we show that *Pf*HtrA2-iKD caused increase in mitochondrial ROS levels and parasite death.

A specific and cell permeable inhibitor of HtrA2 protease, ucf-101, was identified from a high throughput screening of a combinatorial library [42]; the ucf-101 has been used in dissecting the role of HtrA2 in cell survival and cell-death. Since *Pf*HtrA2 was found to be essential for parasite survival, we assessed if protease activity of *Pf*HtrA2 is critical for its functional role. The ucf-101 could effectively inhibit protease activity of recombinant *Pf*HtrA2; however, the ucf-101 showed no effect on growth of parasites, which suggest that the *Pf*HtrA2 protease activity is not essential for parasite survival. Indeed, HtrA2 protein is also suggested to play role in cellular homeostasis through maintenance of mitochondrial proteins in properly folded form [20], whereas the activated HtrA2 protease plays a pro-apoptotic role under cellular stress [14, 43].

It has been shown that the PDZ domain of HtrA2 plays key role in regulating the dual role of HtrA2 in human cells; during normal cell cycle, the PDZ domain of HtrA2 blocks the protease active site and mediates protein-protein interactions, so that a large membrane associated protein complex is formed that acts as a chaperone for protein quality control in the mitochondria [36]. The C-terminal PDZ domain is known to be involved in protein-protein interactions and regulation of the activity of the protease domain[10, 44, 45]. It has been shown that HtrA2 exists as an inactive trimer, where PDZ domains restrict substrate access to the protease domain [17, 46, 47]. Structural studies have also shown that PDZ domain plays the role of regulatory domain and it supervises the proteolytic activity of HtrA2 to avoid undesirable proteolysis [48–50]. However, under cellular stress/apoptotic stimuli, the N-terminal processing and phosphorylation leads to conformational change which removes the inhibitory effect of PDZ from the active site [32, 51]. Indeed, our data confirmed interaction of PDZ-domain with protease-domain of the *Pf*HtrA2, this interaction inhibited the protease activity confirming role of C-terminal PDZ in regulating the *Pf*HtrA2 protease activity.

To understand dynamic role of *Pf*HtrA2, we utilized two cellular stress induction systems: we have earlier shown that proteasome inhibitor MG132 induces ER-stress in the parasite, which leads to apoptosis like cell-death [7]; in another study, we have shown that transient and regulated expression of an active-site mutant of an essential mitochondrial ClpQ protease, *Pf*ClpQ, disrupts this protease system, which causes mitochondrial dysfunction and leads to apoptosis like cell-death [8]. Here we show that the cell -eath induced by either of these cellular stresses is mediated by activation of caspase-like VAD-FMK binding proteases. The activation of caspases is an important step towards the progression of apoptosis like cell-death. Kinetoplastids such as *Plasmodium* and *Trypanosoma* lack conventional caspases; instead, they have meta-caspases (MCAs), which have a C14 domain with a cysteine and histidine caspase-like catalytic dyad [52], in addition, the pan-caspase inhibitor z-VAD-FMK is shown to inhibit cell-death in *P. falciparum* [53, 54] suggesting role of such caspase-like proteases in cell-death. In previous study, it has been shown that VAD-FMK-binding proteases activation is responsible for degradation of a highly conserved metacaspases substrate TSN (Tudor staphylococcal nuclease) in *Plasmodium* [55].

Using these two cellular stress systems we analysed the possible role of *Pf*HtrA2 in activation of Caspase like proteases and subsequent cell-death. We observed a processing event of *Pf*HtrA2, suggesting generation of an activated protease domain under ER or mitochondrial specific stress conditions. In mammalian cells under normal growth conditions the Inhibitor of Apoptosis proteins (IAPs) are known to down-regulate the caspases by either obstructing with their processing or by binding with their active site directly [56]; under apoptotic conditions HtrA2 plays role in removal of IAPs inhibitory effect by directly binding with IAPs through its N-terminal AVPS motif. This active form of HtrA2 also acts like a pro-apoptotic protein by virtue of its protease domain. Sequence analysis of *Pf*HtrA2 showed that the motif AVPS is substituted for the motif ALPS, which is present just at the C-terminus of the protease domain. The presence of IAPs have not been reported in *P. falciparum*, however activation of *Pf*HtrA2 in stressed parasites may suggest its role in regulation/activation of caspase-like proteases. In order to decipher the role of *Pf*HtrA2 in activation of caspase-like proteases, we used a specific inhibitor of HtrA2, ucf-101. Inhibition of *Pf*HtrA2 by ucf-101 effectively reduced activation of the caspase-like protease as well as apoptosis cell-death, as evident by reduction in TUNEL staining, in both ER-stressed and mitochondrial-stressed parasites. These effects were similar as with the pan-caspase inhibitor z-VAD-FMK under both the cellular stress conditions. Overall, these results suggest that *Pf*HtrA2 gets activated as a protease under cellular stress conditions and plays role in the activation of caspase-like proteases which lead to subsequent apoptosis-like cell-death. As described above, it is shown that the activated HtrA2 under apoptotic conditions relocates to cytosol, degrades IAPs and activate caspases [22, 25]; in these systems extra-mitochondrial expression of HtrA2 is also able to induce cell-death [14]. However, under certain cellular stress conditions, the processed HtrA2 remains in the mitochondria to induce pro-apoptotic effect; it is shown that hypoxia induced cellular stress cause loss of membrane potential to promote translocation of HtrA2 into the mitochondrial matrix, where it cleaves protein substrates responsible for maintaining mitochondrial homeostasis and thus promotes cell-death [3, 57, 58]. An important distinction is that in our study the activated *Pf*HtrA2 did not show mobilization into cytosol under cellular stress conditions and stayed in the mitochondria; which may suggest its role in cell death by acting within mitochondria. To further ascertain that the induction of cell-death is not due to any cytosolic translocation of activated *Pf*HtrA2-protease, a transgene corresponding to *Pf*HtrA2-protease was transiently overexpressed in parasite cytosol under degradation-domain based regulation. Since PDZ domain was found to inhibitory to protease activity, the PDZ domain was not included in the transgene. Transient expression of *Pf*HtrA2-protease did not cause any induction of caspase-like protease or cell-death in the parasites. These results suggest that the activated *Pf*HtrA2 plays role in targeting protein substrate in the parasite mitochondria, which disrupts mitochondrial homeostasis and causes activation of apoptosis cascade leading to parasite cell death. Overall, this study highlights the role of *Pf*HtrA2 in mediating parasite organelle homeostasis and cell-death, these results could be helpful to design novel anti-malarial strategy.

## Material and methods

### Parasite culture, plasmid constructs and parasite transfection

*Plasmodium falciparum* 3D7 strain parasites were cultured within human O+ erythrocytes maintaining 4% haematocrit in RPMI 1640 medium (Invitrogen Corp., San Diego, CA, USA) supplemented with Albumax I (0.5%) and hypoxanthine (0.001%) following the standard culture protocols [59, 60]. Parasite culture was synchronized with repeated sorbitol treatment as previously described protocol [61]. To generate *Pf*HtrA2-HA-*glm*S construct for endogenous tagging of *pfhtrA2* gene, the C-terminal fragment of *PfhtrA2* (PF3D7_0812200, 743 bp except the stop codon) was cloned into pHA-glmS vector [62] using the *Bgl*II and *Pst*I restriction enzyme sites. Tightly synchronized ring stage parasites were transfected with 100 µg of *Pf*HtrA2-HA-glmS plasmid by electroporation (310V, 950 μF) [63]. Transfected parasites were selected on WR99210 drug pressure, subsequently subjected to on and off drug cycles to get parasites with plasmid integrated in the main genome and further clonal selection was done to get pure integrant clones by limiting dilution method. Integration of HA-*glm*S at C-terminus was confirmed by PCR using specific primers, as well as by western blot analysis using anti-HA antibody.

To generate constructs for trans-expression of *Pf*HtrA2-protease domain and its mutant form, gene fragment corresponding to 138aa-284aa region of *Pf*HtrA2 was fused with HA-DDD (*Escherichia coli* DHFR degradation domain) tag at its C-terminus was cloned in plasmid vector pSSF2 to generate constructs *Pf*HtrA2-P-DDD. In another construct, same fragment with triple mutations for the active site residues (His154Ala; Asp184Ala; and Ser263Ala) was cloned in similar way to generate the construct *Pf*HtrA2-P(mut)-DDD. The parasites were transfected with each of these constructs separately and selected over WR99210 drug pressure as described above. Expression of fusion proteins [*Pf*HtrA2-P-DDD or *Pf*HtrA2-P(mut)-DDD] was induced in presence of 5μM trimethoprim (TMP) in the culture.

### Conditional knock-down, phenotypic analysis and *in-vitro* growth assays

To assess the effect of *Pf*HtrA2 HA-*glm*S transgenic parasites, we carried out *glm*S mediated conditional knockdown. Tightly synchronized parasite cultures at early ring stage (6-8 hpi) parasites were incubated with glucosamine (2.5 mM or 5 mM) or solvent alone, and allowed to grow for three consecutive cycles. To assess the effect on growth and morphology of parasite, thin smears of parasite culture were made from each well at different time points (0, 24, 48, 72, 96 and 120 h) and stained with Giemsa stain for microscopic analysis. The parasitemia was determined in the next cycle using flow cytometry as well as by counting the parasites in Giemsa-stained smears. Flow cytometry analysis was carried by using FACSCalibur flow cytometer and CellQuestPro software (Becton Dickinson, San Jose, CA, USA). The numbers of ring/trophozoite stage parasites per 5000 RBCs were determined in Giemsa-stained smears and percentage parasitemia [(number of infected erythrocytes/total number of erythrocytes) ×100] was calculated to assess the parasite growth inhibition. Each of the assay was performed three times separately on different days.

### Parasite fractionation and Western blotting

Western blot analysis was carried out to assess expression of *Pf*HtrA2-HA-*glm*S in the transgenic *P. falciparum* blood stage parasites. Briefly, parasites were lysed by 0.15% saponin, the supernatant and washed pellets were separately suspended in Laemmli buffer, boiled, centrifuged, and the supernatant obtained was separated on 12% SDS-PAGE. The fractionated proteins were transferred from gel onto the PVDF membrane (Amersham, Piscataway, NJ, USA) and blocked-in blocking buffer (1 × PBS, 0.1% Tween-20, 5% skimmed milk powder) for 2 h. The blot was washed and incubated for 3 h with primary antibody [Rabbit anti-HtrA2 (1:500); rabbit anti-BiP (1:10000); rat anti-HA (1:1000)] diluted in dilution buffer (1 × PBS, 0.1% Tween-20, and 1% milk powder). Subsequently, the blot was washed and incubated for 1 h with appropriate secondary antibody (anti-rabbit, anti-rat or anti-mouse, 1:20000) conjugated to HRP. Bands were visualized by using ECL detection kit (Amersham) with Biorad ECL Chemidoc imaging system

### Organelle staining, immuno-labelling and fluorescence microscopy

To visualize the mitochondria, the transgenic parasites were stained with MitoTracker Red CMXRos (Invitrogen) at final concentration of 50 nM for 20 mins at 37°C. For immuno-staining, cells were washed with 1× PBS and subsequently fixed in 4% paraformaldehyde. Immuno-labelling was done by incubation of fixed and permeabilized parasites with the primary antibody, anti-HA (1:500) for 3 hours and subsequently with the secondary antibody, Alexa488 goat anti-rat (1:500) for 1 h. The nuclei of parasites were stained using 4′,6-diamidino-2-phenylindole (DAPI, Sigma) with final concentration (2 μg/ ml). The stained parasites were visualized using a Nikon A1 confocal laser scanning microscope and images were analysed using Nikon –NIS element software (version 4.1). The 3D images were constructed by using series of Z-stack images using IMARIS 7.0 (Bitplane Scientific) software.

### Mitochondria membrane potential assay, mitochondrial oxidative stress measurement and analysis of activation of Caspase-like proteases

The mitochondrial membrane potential was assessed by using JC-1 staining dye as described earlier [7]. Infected erythrocytes from parasite cultures in control and experimental sets were incubated with JC-1 (5,50,6,60 -tetrachloro-1,10,3,30 -tetraethylbenzimidazolyl-carbocyanine iodide) at a final concentration of 5 µM, for 30 mins at 37°C. After washing with 1× PBS, the cells were examined by flow cytometry using FACSCalibur flow cytometer and CellQuestPro software (Becton Dickinson, San Jose, CA, USA), using green (488 nm) and red (635 nm) filters. Ratio of JC-1(red)/JC-1(green) was calculated to assess the loss of mitochondrial membrane potential. The JC-1-stained uninfected RBCs were used as background controls.

For mitochondrial oxidative stress measurement, *Pf*HtrA2-HA tagged parasite were grown in the presence and in absence of glucosamine and stained with MitoSOX™ Red (Mitochondrial Superoxide Indicator, Thermo Fisher) at a final concentration of 2μM at 37 °C for 30 min in incomplete media. The stained parasites were washed with 1× PBS. Live MitoSox stained parasite were viewed using a Nikon A1 confocal laser scanning microscope as described above. Images were analysed to determine the mean fluorescence intensities of individual parasites using Nikon -NIS element software (version 4.1).

For Caspase-like cysteine protease activation assay, parasites from treated and control set of culture, were stained with CaspACE FITC-VAD-FMK (fluorescein isothiocyanate-valyl-alanyl-aspartyl-[O-methyl]-fluoromethylketone) *in situ* Marker (Promega, Mannheim, Germany) as per manufacturer’s instructions. The parasites were incubated with 10 µM of CaspACE FITC-VAD-FMK for 30 min at 37°C followed by washing with 1× PBS. The stained samples were analysed by flow cytometry using FACS Calibur flow cytometer and CellQuestPro software (Becton Dickinson) to assess fluorescence staining (Em-525 nm/Ex-488 nm) of infected RBCs. Uninfected RBCs were used as background control.

### Isolation of total DNA, and quantitative real-time PCR

Total genomic DNA was isolated from *Pf*HtrA2-HA-*glm*S parasites grown in presence or absence of glucosamine at different time point. Gene-specific primer sets were designed using Beacon Designer 4.0 software for each organelle and nuclear genome: *tuf*A (apicoplast, *PF*3D7_1348300), *<u>cox</u>*3 (mitochondria, mal_mito_1), as well as for 18S ribosomal RNA (18S rRNA) housekeeping gene as control [64]; details of all the primers are given in Table S1). The amplification reaction contained 10ng template genomic DNA, 2× Maxima SYBR Green qPCR Master Mix (Thermo Scientific) and 10 nM gene specific primers. Real-time PCR was performed in Micro Amp optical 96 well plates in automated ABI Step one Plus Version. Threshold cycle (Ct) values were calculated using SDS 2.4 Software (Applied Biosystem). Standard curves were used to determine genome equivalents of Ct values for respective gene and 18S rRNA gene for each sample. Genome equivalents of each gene was normalized using that of 18S rRNA gene for all the samples; the organelle: nuclear genome ratio was calculated relative to that of control sample as described earlier [9].

### *In vitro* protein-protein interactions assays

*In vitro* protein-protein interaction was assessed by solution binding of the two proteins followed by immuno-pulldown of the protein complex, as described previously [7]. Briefly, 1µg of both *Pf*HtrA2-PDZ and *Pf*HtrA2-protease recombinant proteins were incubated in 100 µl of binding buffer (50mM phosphate buffer pH-7, 75mM NaCl, 2.5mM EDTA pH-8.0, 5mM MgCl_2_, 0.1% NP-40 and 10mM DTT) for 2hr. Subsequently, the complex reaction mixture was incubated for 2hr at 4°C with protein A Sepharose beads, having immobilised anti-HtrA2-PDZ or anti-HtrA2-protease domain antibody. Beads were washed extensively with binding buffer; bound proteins were solubilized in SDS-PAGE buffer and analysed by immunoblotting using anti-HtrA2-PDZ antibody. In control reactions, a non-specific anti-*Pf*HDP antibody were immobilised on sepharose beads in place of the *Pf*HtrA2 antibody.

*In vitro* protein-protein interaction was also analysed with solid surface interaction based technique, where bound proteins were detected by ELISA. Briefly, 96 well microtiter plate Maxi-Sorp micro-titter plates (Nunc International, Nunc, Langenselbold, Germany) was coated with 100 ng of *Pf*HtrA2-PDZ protein in each well. After washing three times with 1× PBS containing 0.05% Tween-20 (PBS-T), the wells were blocked with 1% BSA for 2hr. Recombinant protein *Pf*HtrA2-Protease was added in varying concentration (25-200ng) into *Pf*HtrA2-PDZ coated wells and incubated for 3hr at room temperature. After repeated washing with PBS-T, the wells were sequentially incubated with rabbit anti-HtrA2-protease domain antibody (1:2000) and HRP conjugated anti rabbit antibody (1:3000). After consecutive washes, enzyme reactions were developed with substrate o-phenylenediamine dihydrochloride-H_2_O_2_. The resulting absorbance was measured at 490nm using SYNERGY HTX multi-mode micro plate reader. A non-specific protein (*Pf*HDP) was used in place of *Pf*HtrA2-Protease as the negative control.

### Surface-plasmon resonance (SPR) based analysis of protein-protein interaction

The real time protein-protein interaction of protease and PDZ domain proteins of *Pf*HtrA2was analysed by SPR using Biacore T200 instrument (GE Healthcare, Uppsala, Sweden) as described previously [7]. Briefly, Sensor chip CM5 dextran matrix was activated with a mixture of 1-ethyl-3-3-dimethylaminopropyl)-carbodiimide (EDC) and N-hydroxysuccinimide (NHS). Recombinant protein *Pf*HtrA2-PDZ (100µg/ml in HBS-EP Buffer) was immobilized over the activated surface of the sensor chip flow cell at rate of 10µl/min and 14769 RU was achieved. Another sensor chip flow cell was blocked with 1M ethanolamine-HCl pH 8.5, to use as control. Kinetic binding analysis was done by injecting increasing concentration of *Pf*HtrA2-protease recombinant protein at flow rate of 30 μl/min on immobilized surface. The kinetic parameters were analysed using Biacore evaluation software, version 4.1.1 (GE Healthcare).

### Statistical Analysis

The data were compared using unpaired Student’s t test; the data sets were analysed and the graphical presentations were made using GraphPad Prism ver 5.0.

## Data Availability

All data generated is included in this article and its supplementary information files.

## Competing interests

The authors declare that they have no competing interests

## Acknowledgements

We are grateful to Paul Gilson for providing vector pHAglmS, and Siggi Sato for providing pSSPF2 vector. We thank Rotary blood bank, New Delhi for providing the RBCs. We thank Naresh for his assistance in SPR analysis.

## Funding

The research work in AM’s laboratory is supported by Centre of Excellence grant (BT/COE/34/SP15138/2015) and Flagship Grant (#BT/IC-06/003/91) from the Department of Biotechnology, Govt. of India, and Indo-French Collaborative Research Program Grant (Project 6003-1) by the CEFIPRA. GD and SJ are supported by research fellowships from CSIR, Government of India. VT is supported by BioCARE grant and Az M is supported by research fellowship from Department of Biotechnology, Govt. of India. MA is supported by research fellowship from ICMR, Govt. of India.

## Authors’ Contributions

AM conceived and designed the study; SS, GD, SJ, VT, Az M and MA carried out the experiments; AM, and SA analysed the data; and AM, SS and GD wrote the paper with contributions from all the authors.

